# Decoding images in the mind’s eye: The temporal dynamics of visual imagery

**DOI:** 10.1101/637603

**Authors:** Sophia M. Shatek, Tijl Grootswagers, Amanda K. Robinson, Thomas A. Carlson

## Abstract

Mental imagery is the ability to generate images in the mind in the absence of sensory input. Both perceptual visual processing and internally generated imagery engage large, overlapping networks of brain regions. However, it is unclear whether they are characterized by similar temporal dynamics. Recent magnetoencephalography work has shown that object category information was decodable from brain activity during mental imagery, but the timing was delayed relative to perception. The current study builds on these findings, using electroencephalography to investigate the dynamics of mental imagery. Sixteen participants viewed two images of the Sydney Harbour Bridge and two images of Santa Claus. On each trial, they viewed a sequence of the four images and were asked to imagine one of them, which was cued retroactively by its temporal location in the sequence. Time-resolved multivariate pattern analysis was used to decode the viewed and imagined stimuli. Our results indicate that the dynamics of imagery processes are more variable across, and within, participants compared to perception of physical stimuli. Although category and exemplar information was decodable for viewed stimuli, there were no informative patterns of activity during mental imagery. The current findings suggest stimulus complexity, task design and individual differences may influence the ability to successfully decode imagined images. We discuss the implications of these results for our understanding of the neural processes underlying mental imagery.

## Introduction

Does the Mona Lisa face left or right? A common method of solving this problem is to form an image of the Da Vinci painting in your ‘mind’s eye’. Our ability to imagine scenes and objects can help us solve everyday problems and accomplish day-to-day tasks, such as retracing our steps to find a lost item or navigating from a memorised map. These mentally-generated images are formed in the absence of visual information, and are instead based on short- or long-term memories (Ganis et al., 2003; Kosslyn et al., 2001). Images generated from memory seem anecdotally weaker, or less vivid, than those evoked by sensory input, yet also appear to rely on the visual system (Dijkstra et al., 2018). In line with this, current theories of mental imagery involve common mechanisms for human vision and mental imagery.

Recent work has revealed overlapping neural substrates for visual perception and imagery. Positron emission tomography (PET) and functional magnetic resonance imaging (fMRI) have revealed similar patterns of brain activity during perception and imagery, suggesting computational overlap in the neural systems responsible for each process (Ganis et al., 2004; Kosslyn et al., 1999; Lee et al., 2012; Slotnick et al., 2005). This overlap is particularly clear for areas associated with higher-order abstract visual processing, such as visual association cortex (Albers et al., 2013; Goldenberg et al., 1989; Knauff et al., 2000) and category-selective temporal cortices (Mechelli et al., 2004; Reeder et al., 2015). Overlapping activation is also present in low-level visual areas, despite the absence of visual input during imagery; imagery and visual perception both activate the lateral geniculate nucleus of the thalamus (LGN) (Chen et al., 1998) and primary visual cortex (V1) (Albers et al., 2013; Harrison and Tong, 2009; Pearson et al., 2008). Together, this supports the notion that imagery utilises many of the same mechanisms as visual perception.

Despite overlapping neural activation for vision and imagery, the neural processes are not identical. For example, there is more overlap in higher, anterior regions (i.e., frontal and parietal; Ganis et al., 2004), compared to lower, posterior visual regions (Harrison and Tong, 2009; Lee et al., 2012). There are also task-related differences in imagery such that different imagery tasks show varying degrees of overlap with vision (Ganis et al., 2004; Ishai et al., 2000; Kosslyn and Thompson, 2003). Patients with brain damage also provide evidence for dissociation between imagery and vision. Some patients with occipital or parietal lesions can successfully complete tasks relying on mental imagery, despite significant visual deficits, while others have fully functioning vision but impaired imagery (Bartolomeo et al., 2013; Bridge et al., 2012; Moro et al., 2008; Zago et al., 2010). Therefore, there is some dissociation between vision and imagery despite similar neural processing.

To date, research has focused on understanding the brain networks recruited by a variety of imagery tasks (Fulford et al., 2018; Mechelli et al., 2004), yet we have very little understanding of the temporal dynamics of mental imagery. Although fMRI studies have found correlations between imagery and perception in the later stages of visual processing (Stokes et al., 2011), as well as similar activation patterns between imagery and working memory (Albers et al., 2013), this evidence is limited by the temporal resolution of fMRI. Recent work using MEG has revealed that while similar activation patterns are present in imagery and vision, they occur at a later time and are more diffuse, pointing towards a temporal dissociation between the two seemingly similar processes (Dijkstra et al., 2018).

Multi-Variate Pattern Analysis (MVPA) applied to neuroimaging data can elucidate the information represented in different brain regions (fMRI), and at particular points in time (M/EEG). MVPA offers an advantage in analysing data from mental imagery, as analyses are conducted at an individual-subject level and mental imagery ability is understood to vary significantly between people (e.g., Cui et al., 2007). MVPA is also more sensitive to variation across fine-grained patterns, and provides a powerful framework for the detection of content-specific information (Grootswagers et al., 2017; Haynes, 2015). This is particularly advantageous for imagery signals that are likely to be weaker than visual input (Naselaris et al., 2015). One recent study found that the category of imagined images (faces and houses) was decodable from MEG recordings, albeit later than viewed images (Dijkstra et al., 2018). However, decoding of individual exemplars was poor, indicating a dissociation between low- and high-level imagery processes.

Here, we examined how the neural representation of mental images develops and changes over time. Participants imagined one of four previously learned pictures: two faces and two places. Each image was visually dissimilar to the other within the category, while maintaining clear category divisions. Neural responses were measured using EEG while participants viewed the experimental images, imagined the images, and viewed fast streams of semantically related images (i.e., other faces and places). We expected that category information would be decodable from the EEG data during mental imagery (Dijkstra et al., 2018), that it would be broadly generalisable across the imagery period, and delayed relative to vision. We also predicted that exemplars within each category would be distinguishable (i.e., successful within-category decoding). We found that the dynamics of imagery processes are more variable across, and within, participants compared to perception of physical stimuli. Although category and exemplar information was decodable for viewed stimuli, there were no informative patterns of activity during mental imagery.

## Materials and Methods

### Experimental structure

At the start of the session, participants completed the Vividness of Visual Imagery Questionnaire (VVIQ) (Marks, 1973). They were then informed of the task instructions and completed 24 imagery task training trials. The experiment itself consisted of four blocks that were completed while EEG was measured. In each block, participants passively viewed five rapid streams of images (Pattern Estimator), followed by a series of imagery trials. Each imagery trial consisted of a four-image sequence (Seen images), after which participants were cued to imagine one of those stimuli (Imagery).

### Participants

We recruited 16 right-handed subjects (11 male), of mean age 23 (SD= 5.58, range 18-39), with normal or corrected-to-normal vision and no history of psychiatric or neurological disorders. The experiment was approved by the Human Ethics Committee of the University of Sydney. Written, informed consent was obtained from all participants.

### Behavioural data

To measure individual variation in vividness, we administered a modified VVIQ (Marks, 1973) prior to EEG set-up. The VVIQ measures subjective perception of the strength of an individual’s mental imagery. Participants were asked to imagine 16 scenarios, and rated each for vividness on a five-point Likert-like scale. A reversed scoring system was used to decrease confusion. Participants rated each item from 1 (*“No image at all, you only ‘know’ that you are thinking of an object”)*to 5 (“*Perfectly clear and as vivid as normal vision”*). All questions were completed twice, once with open eyes and once with closed eyes. A final summed score between 32 and 160 was calculated for each subject; higher scores indicate greater vividness.

### Apparatus and Stimuli

Four stimuli were used in this experiment: two images of Santa and two images of the Sydney Harbour Bridge. The inclusion of two exemplars per category allowed us to disentangle whether participants are thinking of the concept (i.e., Santa, Sydney Harbour Bridge) or generating a specific image. These stimuli also fit into distinct face/place categories, which have been shown to evoke robustly distinct patterns of neural activity (Haxby et al., 2001; Kanwisher et al., 1997).

All stimuli were displayed on a 1920 x 1080 pixel Asus monitor on a grey background. Participants viewed stimuli at approximately 57cm, such that all stimuli subtended approximately 4.1 degrees of visual angle (including a 0.15 degree black border). Responses were made using a mouse with the right hand. A grey fixation cross was superimposed on all stimuli, with horizontal and vertical arms subtending approximately 0.6 degrees of visual angle. Experimental presentations were coded in MATLAB using extensions from the PsychoPhysics Toolbox (Brainard, 1997; Kleiner et al., 2007; Pelli, 1997).

### Imagery sequence

Each imagery sequence began with a fixation cross in the centre of the screen for 1000 milliseconds. The four stimuli were displayed sequentially in the centre of the screen, within a black border. Each was displayed for 1500 milliseconds each, in a pseudo-random order. Targets were counterbalanced such that each block contained all 24 possible sequences of the four stimuli. For each sequence, a different target was selected in each block. Target allocation in each block was also randomised. This counterbalancing meant each image appeared in each temporal position as a target equally often.

The fourth stimulus was followed by a 1000ms fixation cross, then a numerical cue appeared (1-4). This cue referred to the target’s position in the stream; for example, ‘3’ indicated the target was the third image in the stream. Participants were instructed to click the mouse once they had identified the target and were mentally “projecting an image into the square”. Upon clicking, the number was replaced with a dark grey fixation cross and the frame was filled light grey. This ‘imagery’ screen was displayed for 3000ms before automatically advancing to a response screen. On the response screen, participants were shown the four stimuli and horizontal mirror images of these stimuli. They used a mouse to select which of these images they were imagining. Mirror images were used as distractors because they are semantically identical but visually different, to determine if participants were using a semantic strategy rather than an imagery-based strategy. Horizontal positioning changed across blocks (stimulus identity), and vertical positioning was randomised every trial (mirror images/stimulus) such that for some trials the mirror image was in the top row, and some in the bottom row. This randomisation aimed to reduce predictability in responses.

### Training

Participants completed a block of 24 practice trials of the imagery sequence before EEG recording. We expected these training trials to give participants the opportunity to learn task structure and observe more details about the images to facilitate vivid imagery. Training trials were similar to experimental trials. The first 12 trials contained typed instructions on how to identify the target, and went straight to the response screen after the cue, with no imagery component. On incorrect responses, participants were shown the correct image. The second 12 trials mimicked experimental trials, with the addition of typed instructions and feedback. Participants were given the option to repeat the training, and two did so.

### Pattern Estimator

We also included a pattern estimator at the beginning of each to investigate the degree of generalisation across semantic category. These images were semantically similar to the critical experimental stimuli. Participants passively viewed a rapid stream containing the four stimuli from the imagery sequence, as well as horizontally flipped, inverted and blurred versions of these images. It also included other images of the Sydney Harbour Bridge and Santa, other bridges and other people. Each block began with five short streams of 56 images, displayed for 200ms each. Every stream contained all 56 images in a random order, and lasted for 11.2 seconds. Participants could pause between streams and elected to advance when they were ready.

## Data recording and processing

### EEG recording

EEG data were continuously recorded at 1000Hz using a 64-channel Brain Products (GmbH, Herrsching, Germany) ActiCAP system with active electrodes. Electrode locations corresponded to the modified 10-10 international system for electrode placement (Oostenveld and Praamstra, 2001), with the online reference at Cz. Electrolyte gel kept impedances below 10kΩ.

### Pre-processing EEG

EEG pre-processing was completed offline using EEGLAB (Delorme and Makeig, 2004) and ERPLAB (Lopez-Calderon and Luck, 2014). The data were minimally pre-processed. Data were down-sampled to 250Hz to reduce computational load, then filtered using a 0.1Hz high-pass filter, and a 100Hz low-pass filter. Line noise at 50Hz was removed using the CleanLine function in EEGLAB. Four types of epochs were created: Pattern Estimator, Vision, Cue-Locked Imagined and Response-Locked Imagined. Each epoch included 300ms before to 1500ms after stimulus onset. Pattern Estimator epochs were from the fast stream at the beginning of each block, and Vision epochs were from the four images displayed in each experimental trial. Cue-locked Imagined epochs were centred around presentation of the numerical cue designating the target. Response-Locked Imagined epochs were centred around participants’ mouse click to begin imagery. Although the period between cue and response was variable across trials (Supplementary Fig S2), we expected the period immediately following the cue to provide insight into the initial stages of imagery generation.

### Decoding analysis

All EEG analyses were performed using time-resolved decoding methods, custom-written using CoSMoMVPA functions in MATLAB (Oosterhof et al., 2016). For all decoding analyses, a regularised linear discriminant classifier (as implemented in CoSMoMVPA) was trained to differentiate brain patterns evoked by each image or category of images.

For category decoding, a classifier was trained to distinguish images of Santa from images of the Sydney Harbour Bridge for recordings from the same type (i.e., a classifier trained on data from the Pattern Estimator was tested on another independent portion of the Pattern Estimator data). To determine if exemplars were also uniquely represented, a classifier was trained to distinguish between the two exemplars within each category (e.g., decode the two Santa images). Classifiers were trained and tested for each time point using a 12ms sliding time window (three time points).

To analyse data from the Pattern Estimator and Vision epochs, each presentation sequence was treated as independent. We used a leave-one-trial-out cross-validation approach, where Vision trials were composed of the four stimuli in each imagery sequence and Pattern Estimator trials were composed of a single sequence containing all 56 semantically relevant images. Imagined stimuli were analysed using a leave-two-out cross-validation approach, which took each imagery epoch as independent and left one exemplar of each category (one Santa and one Sydney Harbour Bridge) in the test set. Cross-decoding analyses were conducted using split-half cross-validation, where a classifier was trained on one trial type and tested on another trial type (e.g., train on all Vision trials and test on all Cue-Locked Imagined trials). To investigate the possibility of similar processes occurring in vision and imagery at different times, we used temporal generalisation methods (King and Dehaene, 2014), in which the trained classifier for a single time point is applied to every time point in a second set of data.

To compute statistical probability for all within-type, cross-decoding and time generalisation analyses, we used the Monte Carlo Cluster Statistics function in the CoSMoMVPA toolbox (Maris and Oostenveld, 2007; Smith and Nichols, 2009; Stelzer et al., 2013). These statistics yield a corrected p-value that represents the chance that the decoding accuracy could have come from a null distribution formed from 10,000 iterations (North et al., 2002). These p-values were thresholded at *p*_corrected_ < .05 for significance.

## Results

In this experiment, participants viewed rapid streams of images (Pattern Estimator), and series of imagery trials. In imagery trials, participants were presented with a sequence of four images (Vision) and then were cued to imagine one of the images (Imagery). We trained and tested multivariate classifiers to decode exemplar and category of the object in all three conditions, as well as tested the generalisation performance of classifiers between vision and imagery trials.

### Behavioural results

#### Vividness of Visual Imagery Questionnaire

The VVIQ was scored out of 160, a sum of responses to each of the 16 questions on a five-point scale. The VVIQ was given to participants both with eyes open and closed (Marks, 1973). The average overall score was 113 (*SD* = 15.93, range 82-150), similar to previously reported means (Amedi et al., 2005; Crawford, 1982; Fulford et al., 2018). Responses with eyes open (*M* = 56.44, *SD* = 8.54) were very similar to eyes closed (*M* = 57.69, *SD* = 10.28). The distribution of overall scores is shown in Supplementary Figure S1.

#### Target identification

To verify if participants were able to identify the target for imagery trials correctly, we examined their behavioural responses after each imagery sequence. Participants were able to accurately identify the target, with an average overall accuracy of 92% (*SD* = 4.40). Of the trials which were errors, most participants chose one of the four original images (67% of errors). Approximately a third of incorrect responses were to the flipped version of the target. This suggests participants successfully learned the basic characteristics of the target images and were not simply relying on a mnemonic strategy to complete the task. The mean response time from cue to imagery was 3.21 seconds (*SD* = 1.86) and the most frequent response time was between 1.5 and 2 seconds (Supplementary Fig S2).

### EEG results

#### Significant decoding of image category and exemplars for seen images on imagery trials

To test whether category information was represented in visually displayed images, we trained and tested a classifier on the images seen during experimental trials (Vision). Category decoding was continuously above chance (*p*s < .05) after 88ms (Fig 1), indicating patterns of brain activity for Santas and Sydney Harbour Bridges were distinguishable from this point. This above-chance decoding was sustained for the entire time the image was displayed. Continuous above-chance decoding began for both Santas and Sydney Harbour Bridges at 96ms. Peak accuracy occurred at 132ms for Santas, 124ms for Sydney Harbour Bridges and at 196ms for category decoding.

**Figure 1.**
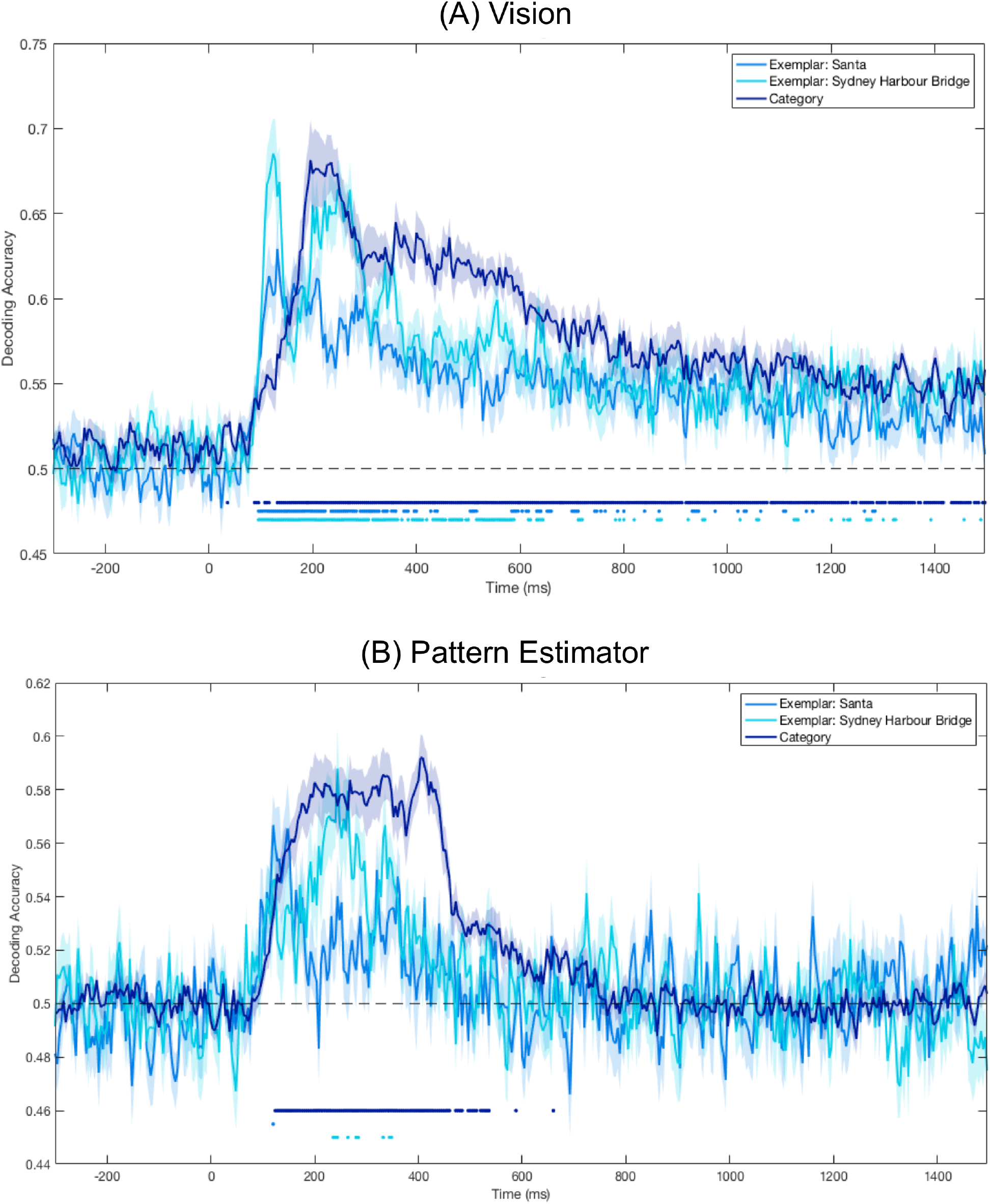
Mean decoding accuracy for Vision (A) and Pattern Estimator (B) images. Dots below plots indicate time points at which decoding was significantly above chance (*p* < .05). Shaded areas represent the standard error of the mean across subjects. (A) Decoding category and exemplar identity from the four target images presented in the experimental trials. (B) Decoding category and exemplar identity from the 56 images presented in the fast streams at the beginning of each block; category decoding was based on all images in the stream classified by either face or place, and exemplar decoding was based only on the targets and modified targets

### Significant category decoding in Pattern Estimator

To create a category classification model for imagery, we looked at patterns of brain activity while participants were viewing images in the fast stream (Pattern Estimator). All images were labelled according to super-ordinate categories of ‘face’ or ‘place’. To assess the model’s utility, we cross-validated it on the Pattern Estimator trials. There was sustained above-chance category decoding from 124ms after stimulus onset until approximately 535ms after stimulus onset (Fig 1). The classifier was also able to distinguish between the two Sydney Harbour Bridge targets at several discrete time points between 236ms and 348ms after stimulus onset. There was no continuous above-chance decoding for Santas. Category decoding peaked at 404ms after stimulus onset, at 244ms for Sydney Harbour Bridges, and at 120ms for Santas.

### No significant decoding for imagery

To determine if category or exemplar information was decodable from imagined data, we trained and tested a classifier on the Cue- and Response-Locked Imagined epochs (Fig 2). Brain areas activated during imagery are known to vary between individuals (Cui et al., 2007), so we looked at imagery decoding on an individual subject basis. For each subject, we ran a permutation test in which the decoding procedure was run 1000 times, with category labels randomly assigned to the epochs. A *p*-value was calculated for each time point, based on the number of permutations with a greater decoding accuracy than the correct label decoding. We used the False Discovery Rate to correct for multiple comparisons. This test was conducted on both Response- and Cue-Locked epochs, and we found decoding was not significantly above chance for any individual at any time point for either Cue- or Response-Locked data (*p*s > .05).

**Figure 2.**
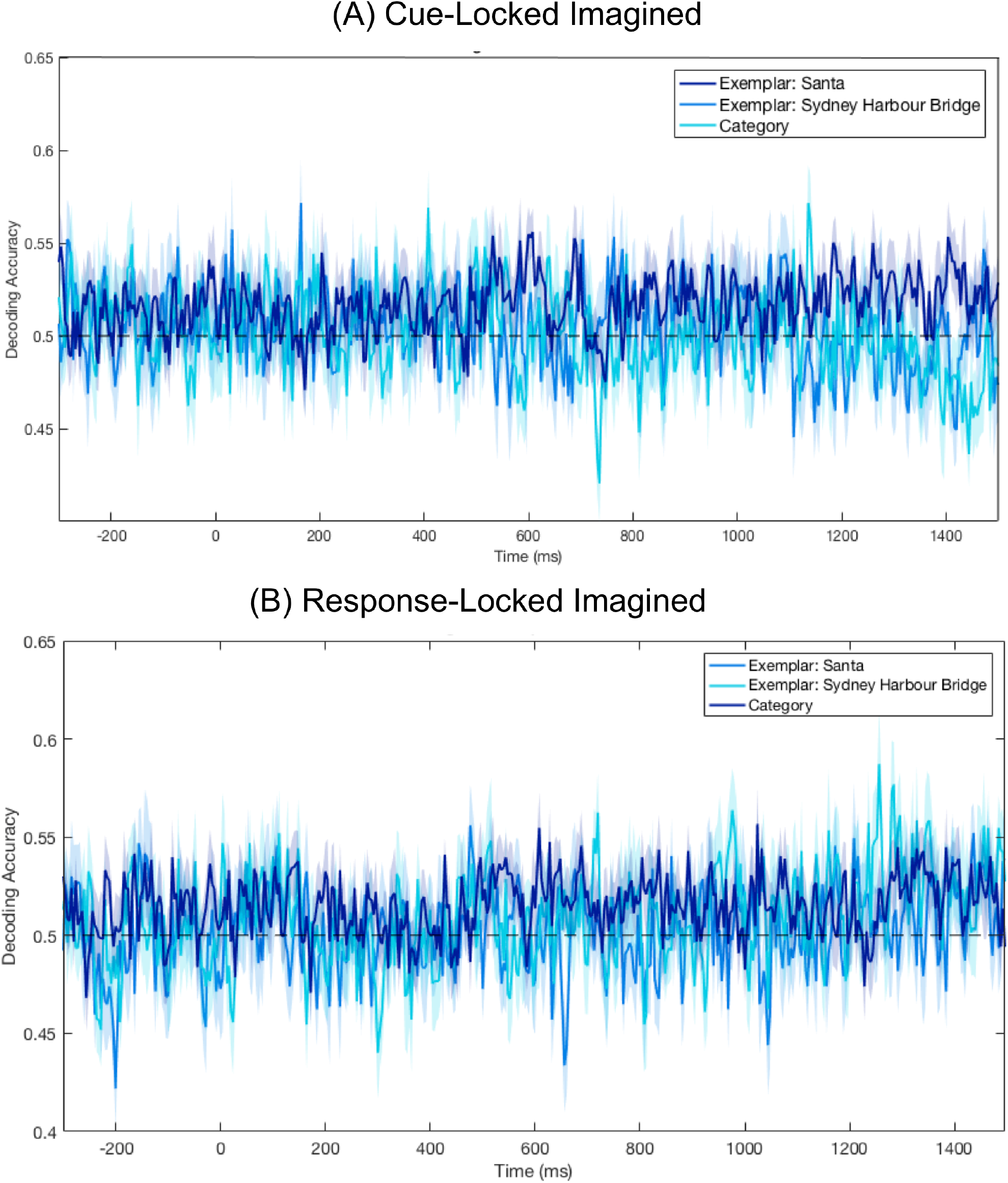
Mean decoding accuracy for Cue-Locked and Response-Locked Imagined epochs. The absence of dots below the plots indicates there were no points at which decoding was significantly above chance (*p*s>.05). Shaded areas represent the standard error of the mean across subjects. **(A)** Decoding accuracy centred on when participants click to advance to the imagining period. **(B)** Decoding accuracy centred on presentation of the numerical cue indicating the location of the target in the preceding stream.

To test whether there was any representational overlap in imagery and vision, we ran a cross-decoding analysis. We ran all pairwise combinations of vision and imagery; a classifier trained to distinguish Santas from Harbour Bridges in the viewed stimuli (Pattern Estimator or Vision epochs) and was tested on imagery periods (Cue-Locked or Response-Locked). There were no significant periods of overlap for any cross-decoding involving imagined trials (*p*s > .05).

It could be that the processes in vision and imagery engage overlapping representations but at different times. To test this, we conducted a time generalisation analysis (King and Dehaene, 2014). A classifier was trained on visual data (Pattern Estimator or Vision epochs) at each time point, and then tested on imagined data (Cue- and Response-Locked) at every possible time point. There was no time point where decoding was significantly above chance for any combination of training and testing (all *p*s > .05), indicating there was no point where the patterns of brain activity during perceptually processed stimuli were present during imagery.

### Differences in vividness did not affect decoding accuracy

Another possibility is that people with greater capacity for imagery have more decodable imagery representations. To investigate the effects of subjective imagery vividness on decoding accuracy, we grouped the participants as ‘high’ or ‘low’ imagery vividness based on a median split of their ‘eyes-open’ scores in the VVIQ. Two participants had the median score and were excluded from further analysis. We used the eyes-open score because it was the most relevant for the task at hand, and makes our results comparable to prior MEG research (Dijkstra et al., 2018), where only the eyes-open section was used. To see if there were any significant differences between the groups in any of the previously described analyses, we conducted a random-effects Monte Carlo statistic with 10,000 iterations to find where differences between the groups were significantly greater than zero. There was only one isolated point of significant differences between the two conditions, at 1484ms, when the classifier was trained on Pattern Estimator data and tested on Response-Locked Imagery.

## Discussion

The current study used time-series decoding to capture the precise temporal fluctuations underlying mental imagery. Based on prior MEG evidence showing the category and identity of imagined objects can be decoded, we expected successful category and exemplar decoding from imagery. However, contrary to our predictions, we were unable to detect any systematic representations of category or exemplar information during imagery. Based on previous evidence that imagery recruits similar neural networks to vision (Ganis et al., 2004), we also anticipated overlapping patterns of neural activity when participants were viewing and imagining the same image. Although we were able to decode stimulus category and identity from visually processed stimuli, there were no time points where neural representations of vision and imagery were overlapping. Finally, we considered whether individual subject results might vary on the basis of imagery vividness, and found no systematic differences between subjects reporting high and low vividness. Overall, our findings demonstrate the variability of imagery processes within subjects over time, and suggest stimulus- and design-related factors may influence the chances of successfully decoding mental imagery.

To compare the overlap between imagery and visual processing, we first defined the temporal dynamics of visual processing for the images in this experiment. For stimuli presented as part of the imagery sequence (Vision), image category was predictable from approximately 100ms after stimulus presentation until offset 1400ms later. Exemplar decoding was also significant from 100ms, albeit for less continuous time than category decoding, reflecting well-established evidence that both categories and exemplars evoke distinct patterns of brain activity (Carlson et al., 2013). For the Pattern Estimator, category decoding was significantly higher than chance from 100ms until approximately 500ms after stimulus onset. This extended period of decoding after stimulus offset supports recent evidence that multiple representations can co-exist in the brain (Grootswagers et al., 2019; Marti and Dehaene, 2017).

In both visual conditions, exemplar decoding peaked earlier than category decoding. This reflects well-established evidence of increasing abstraction along the ventral visual pathway (Carlson et al., 2013; Contini et al., 2017). It also appears that decoding accuracy for Sydney Harbour Bridges is higher than for Santas, for both visual conditions (Vision and Pattern Estimator), though this pattern is less defined for the Pattern Estimator stimuli because of the low numbers of training and testing stimuli (4 of each exemplar per stream). Informal questioning of participants post-experiment suggested many participants found the Sydney Harbour Bridge images easier to imagine because of the distinct lines forming the arches and underside of the bridge.

When the classifier trained on the visual stimuli was tested on imagery, there were no time points where the signal was sufficiently similar to accurately predict image category or identity. To investigate the possibility that the processes were not temporally aligned, we conducted a temporal generalisation analysis. There were no regular patterns of activity at the group level, indicating there was no overlap in representations at any point in the imagery period. Based on evidence that areas of activation during imagery vary across people (e.g., Cui et al., 2007), we examined results on the individual level. Patterns of individual decoding accuracy varied dramatically between subjects. Neither category nor exemplar decoding was significant at any time point for any individual. At face value, these results seem inconsistent with prior findings by Dijkstra and colleagues (Dijkstra et al., 2018). These differences primarily point to the difficulties of studying visual mental imagery, and the specific methodological characteristics required to obtain significant imagery decoding.

Several factors may have impacted our capacity to decode imagined mental representations. For example, the increased number of channels in MEG compared to EEG provides better signal to noise ratio and greater likelihood of detecting an effect (Cichy and Pantazis, 2017). An additional consideration is that individual variability in image generation would reduce the sensitivity of population statistics. Moreover, the temporal variability in an individual’s capacity to generate a mental image would further reduce individual effect sizes.

Another potential explanation for our non-significant imagery decoding is the unavailability of non-imagery based strategies. Previous imagery experiments using a retro-cue design, in which participants identify the imagery target based on a cue presented immediately following a sequence of images, have found significant imagery decoding using only two stimuli (e.g., Dijkstra et al., 2017; Dijkstra et al., 2018; Harrison and Tong, 2009). However, with only two classes of stimuli, participants can effectively complete the task without imagery. For example, participants could perform the retro-cue house-face task used in Dijkstra and colleagues’ research (Dijkstra et al., 2018) by recalling a label for each image as it is presented (e.g., ‘house-face’), and mentally repeating this order after cue presentation. After identifying the target, subjects could simply continue to think of the relevant label. This pattern of thought is likely to be sufficiently similar during perception and imagery to be identified by the classifier as a reliable difference between the categories, leading to accurate decoding of patterns of brain activity based on semantic labels instead of imagery.

We designed our experiment to test if this was the case by including a superordinate category distinction with two exemplars in each category. We obtained response data after every trial with flipped images as distractors to test whether participants were using an imagery-based strategy. If participants were using a purely semantic label-based strategy, we would expect a similar number of responses for flipped and target images. However, only 0.33% of all responses were the flipped version of the target. These response patterns clearly show participants in our experiment were aware of the visual elements of the images rather than solely the semantic label. Due to the fundamentally introspective nature of mental imagery, there is no way to determine if participants are genuinely completing the imagery portion of the task. However, these response patterns point strongly to the use of an imagery-based strategy. Future experiments with similar hierarchical structure and more subtly modified response options (e.g. deleting or rotating a single element of the image, or changing colours of elements of the target images) could help determine whether this is a plausible theoretical explanation for our results.

Generation of mental imagery requires activation of complex, distributed systems (Ganis et al., 2004). Higher stimulus complexity increases the number of details that need to be recalled from memory. It therefore seems likely that the neural processes involved in viewing a static image are more temporally consistent than generating an image from memory, which is unlikely to follow a millisecond-aligned time-locked process. This is particularly apparent for complex stimuli which require more details, stored in potentially disparate locations, to generate vivid imagery. This same temporal blurring between trials from temporally misaligned processes is present in other prior studies (Dijkstra et al., 2018), as it is somewhat inherent to the temporal specificity that decoding of time-series data provides.

Most previous experiments using complex visual scenes as imagery targets use an extensive training period prior to the study, relying on long-term memories of targets for imagery (Naselaris et al., 2015). Although our participants completed a training period prior to EEG recording, slightly longer than those in Dijkstra and colleagues’ MEG study, it is possible (Dijkstra et al., 2018) that participants might have experienced more vivid imagery if they had more exposure to the experimental images. Intuitively, it seems easier to imagine a highly familiar object such as an apple rather than a scene of Sydney Harbour because there are fewer details required to create an accurate representation. Mental images that are less vivid or less detailed are likely to generate weaker neural activation (Dijkstra et al., 2017) and are less likely to fully resemble the details that are processed during vision. If the patterns are less distinct, a classifier is less likely to be able to identify reliable patterns of brain activity on which to base categorisation. To determine the effects of memory on imagery vividness and reliability, future study could compare the current results to a similar paradigm where subjects have extensive training prior to recording (e.g., participants are extensively questioned about characteristics of the image, or have to draw the main aspects to show awareness of details in the image).

As highlighted in recent research (Dijkstra et al., 2019), individual differences in imagery generate increased variation between individuals. For example, differences in visual working memory capacity, personal decision-making boundary, and memory strategy may have increased variation between participants. Individuals who report stronger imagery ability tend to use an imagery-based strategy on visual working memory tasks (Pearson et al., 2015). Features of both working memory and long-term memory (e.g. meaningfulness, familiarity) affect ratings of imagery vividness (Baddeley and Andrade, 2000). These factors might also influence variability within a participant… changes over the course of the experiment, increasing experience with images, etc, could influence temporal variability from trial-to-trial.

Other individual differences, such as personal decision strategies vary across individuals. We may have captured a slightly different stage of imagery, as it is likely each person based the timing of their mouse clicks on a different threshold criterion for the point at which they had begun to imagine. Different strategies for identifying the target may have directed the focus of imagery. When asked informally at the conclusion of the experiment, all participants could explicitly describe their strategy for identifying the target. Most participants assigned a label to each image and mentally repeated these to remember the image order. The majority of strategies relied on structural characteristics, for example, “fat, tall, under, above”. Several participants also reported a direction-based strategy, for example, “top, bottom, centre, side” or “straight, side, face, body”, indicating the orientation of the main object in the image. Though there is no reliable way to compare decoding accuracy based on strategy, different strategies may direct focus on different aspects of the complex images (e.g. thinking of ‘face’ might make facial features salient, compared to labelling the same image as ‘fat’, drawing focus to body shape). These differences in strategy present another potential source of variation between subjects.

It is clear that capability to decode visual mental imagery is influenced by several factors, including vividness, memory and stimulus complexity. These factors do not affect imagery in isolation; they are inherently related. Better memory for the details of an image is likely to increase vividness. The number of details remembered by an individual is influenced by their memory capacity, but also by the complexity of the stimulus and the number of details necessary to generate a vivid image. All these factors create variation in the processes used to generate mental imagery, across both people and time (Borst and Kosslyn, 2010; Dijkstra et al., 2018). The potential for MVPA techniques to analyse data at the individual level provides insight into the variation across subjects, and highlights the need for future studies to consider patterns of data at an individual level to maximise the chances of obtaining clear signals from imagery.

## Conclusion

In this study, we investigated how neural representations of mental imagery change over time. Our results suggest successful category decoding in earlier studies may be a result of better signal to noise ratio from a variety of factors, including individual variation. Variety in response times, imagery strategy and ability, in addition to fewer recording sensors may have reduced our power to find systematic patterns of neural activity during imagery. Furthermore, the interactions between stimulus complexity, working memory, and imagery vividness may have increased this variation between individuals. Our results raise many questions for further investigation and demonstrate both the challenges and advantages associated with time-series decoding for EEG in investigating the introspective processes underlying mental imagery.

## Supplementary Materials

### Author Contributions

Conceptualization, S.M.S.; methodology, S.M.S., T.A.C., A.K.R, T.G.; formal analysis, S.M.S., T.G.; investigation, S.M.S.; writing—original draft preparation, S.M.S.; writing—review and editing, S.S., T.A.C., T.G., A.K.R.; supervision, T.A.C., T.G., A.K.R.; project administration, T.A.C.; funding acquisition, T.A.C.

### Funding

This research was supported by an Australian Research Council Future Fellowship (FT120100816) and an Australian Research Council Discovery Project (DP160101300) awarded to T.A.C. The authors acknowledge the University of Sydney HPC service for providing High Performance Computing resources. The authors declare no competing financial interests.

### Conflicts of Interest

The authors declare no conflict of interest. The funders had no role in the design of the study; in the collection, analyses, or interpretation of data; in the writing of the manuscript, or in the decision to publish the results.

## Supplementary data

**Figure S1.**
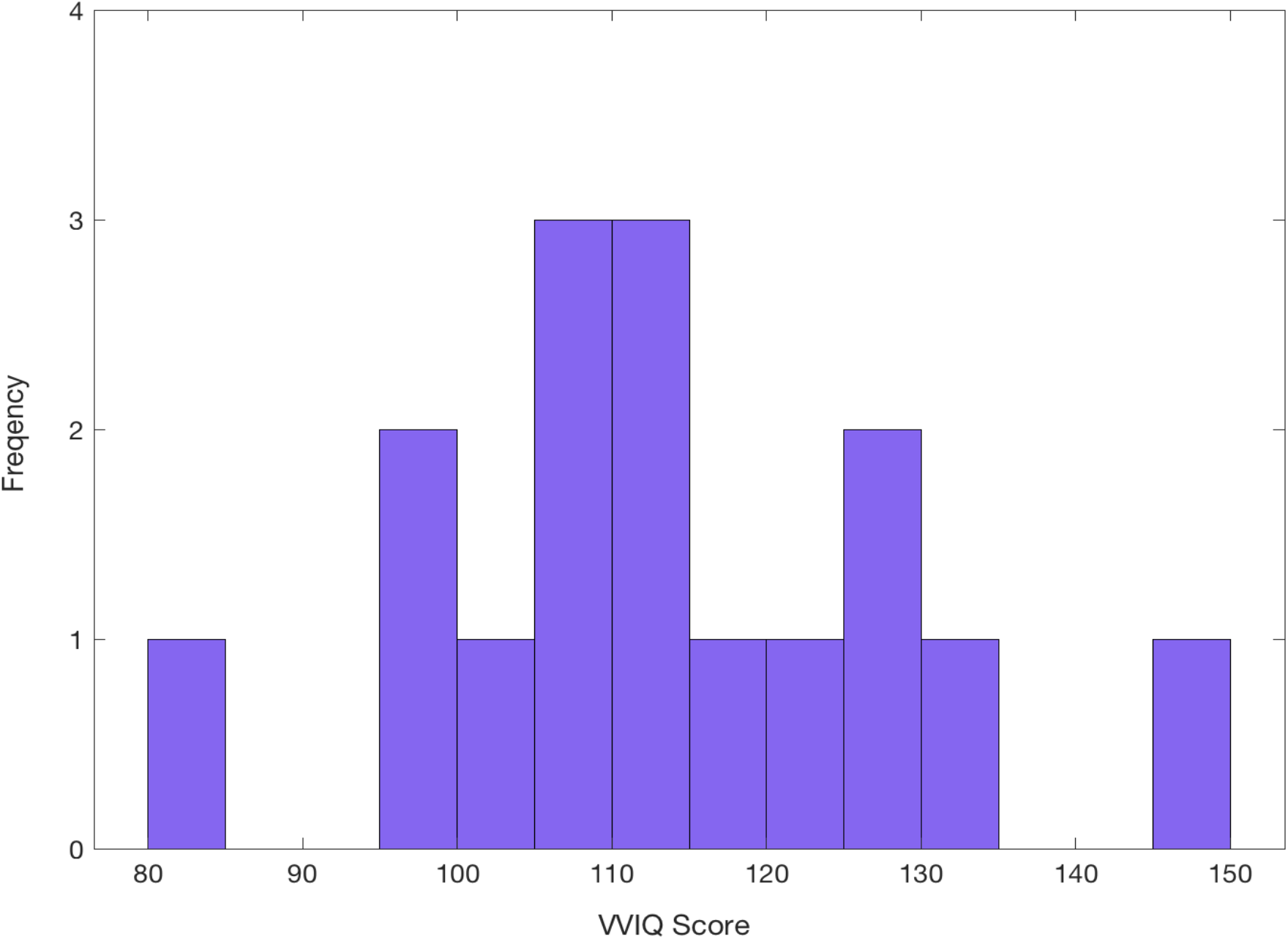
Frequency Distribution of scores in the Vividness of Visual Imagery Questionnaire overall scores. Scores are calculated out of a possible 160 by summing responses to each question completed with the eyes open and with the eyes closed.

**Figure S2.**
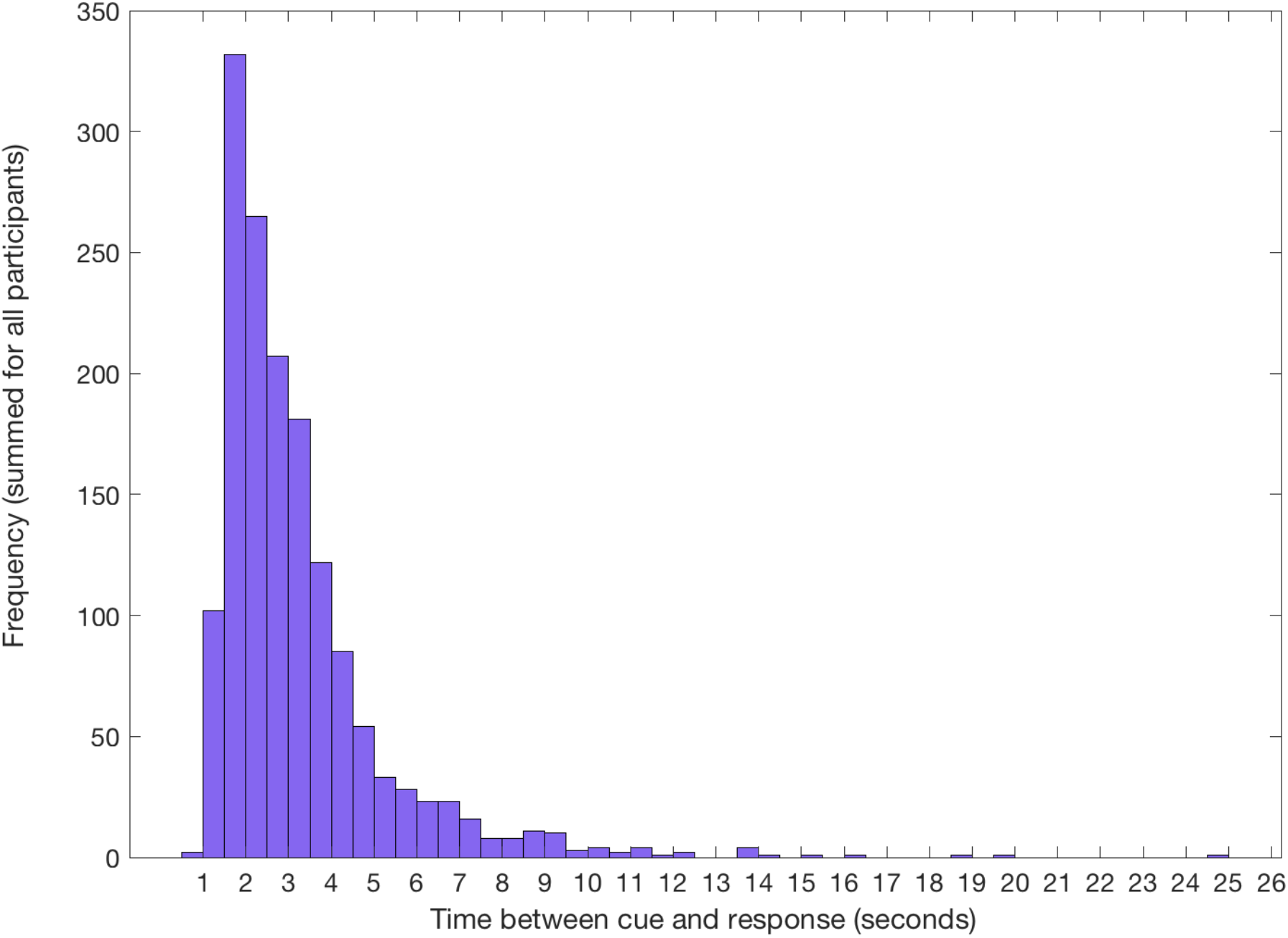
Frequency of response times from cue to imagery across all participants. Response time is taken from the onset of the numerical cue indicating the location of the target in the stream, until the participant voluntarily clicked the mouse. During this period, participants identified the correct target and began to imagine it on the screen.

